# AB-Free Kava and its Kavalactones Suppress Cigarette Smoke- and Lipopolysaccharide-Induced Lung Inflammation: Efficacy and Mechanisms

**DOI:** 10.64898/2026.08.21.746215

**Authors:** Gujie Xu, Tengfei Bian, Breanne Freeman, Yifan Wang, Allison Lynch, Chandra K. Maharjan, Tyler H. Montgomery, Leah R. Reznikov, Adriaan W. Bruijnzee, Weizhou Zhang, Chengguo Xing

**Affiliations:** Department of Medicinal Chemistry and Center for Natural Products, Drug Discovery, and Development, College of Pharmacy, University of Florida, Gainesville, FL 32610, USA; Department of Pathology, Immunology and Laboratory Medicine, College of Medicine, University of Florida, Gainesville, FL 32610, USA; Department of Physiological Sciences, College of Veterinary Medicine, University of Florida, Gainesville, Florida 32610, United States; Department of Psychiatry, College of Medicine, University of Florida, Gainesville, Florida 32610, United States

**Keywords:** AB-free kava, kavalactone, cigarette smoke, LPS, inflammation, IL-6, TNF-α, PGE₂, COX-2, PKA, CREB

## Abstract

Cigarette smoke-induced lung inflammation is a central driver of pulmonary diseases. The limited efficacy of current anti-inflammatory agents underscores the need for structurally novel therapeutics with distinct mechanisms. We recently demonstrated that AB-free kava, a flavokavains A/B-depleted formulation from *Piper methysticum* containing six major kavalactones, effectively suppresses cigarette smoke–induced lung inflammation in mice. This study aims to identify the bioactive constituent(s) and elucidate underlying mechanisms. These kavalactones revealed a clear structure–activity relationship in suppressing lipopolysaccharide (LPS)-stimulated prostaglandin E₂ (PGE₂) production in macrophages with desmethoxyyangonin (DMY) as the most potent kavalactone whereas dihydrokavain (DHK, a structurally similar analog) with minimal activity. DMY also effectively reduced LPS-induced interleukin-6 (IL-6) and tumor necrosis factor alpha (TNF-α) production while DHK was ineffective. Mechanistically, DMY, but not DHK, attenuated COX-2 induction and reduced phosphorylation of cAMP response element–binding protein (CREB). Pharmacological inhibition of protein kinase A (PKA) similarly reduced p-CREB, COX-2 and PGE_2_, supporting a PKA-dependent CREB/COX-2 signaling in mediating PGE₂ suppression while these effects were independent of nuclear factor kappa B (NF-κB) and activator protein 1 (AP-1) signaling. Similar results were observed for DMY and DHK in attenuating cigarette smoke condensate-induced proinflammatory pathways and PGE_2_ production. Consistently, DMY demonstrated significant *in vivo* efficacy in suppressing cigarette smoke– induced lung inflammation while DHK was not effective. Interestingly, dihydromethysticin (DHM) demonstrated the greatest *in vivo* anti-inflammatory efficacy, although it only exhibited moderate *in vitro* potency, likely due to its superior bioavailability over DMY. Concordantly, cigarette smoke exposure elevated p-CREB and COX-2 expressions in mouse lungs, which were attenuated by AB-free kava and its bioactive kavalactones with the extent of suppression correlating with their *in vivo* anti-inflammatory efficacy. DHM effectively suppressed LPS-induced neutrophil accumulation in mouse lungs as well. Collectively, these studies identify bioactive kavalactones in AB-free kava that suppress cigarette smoke- and LPS-induced lung inflammation through the modulation of the PKA/CREB/COX-2 signaling axis, providing a foundation for developing structurally distinct anti-inflammatory agents, particularly targeting smoke-induced inflammation and associated pulmonary diseases.

**Graphic Abstract:** 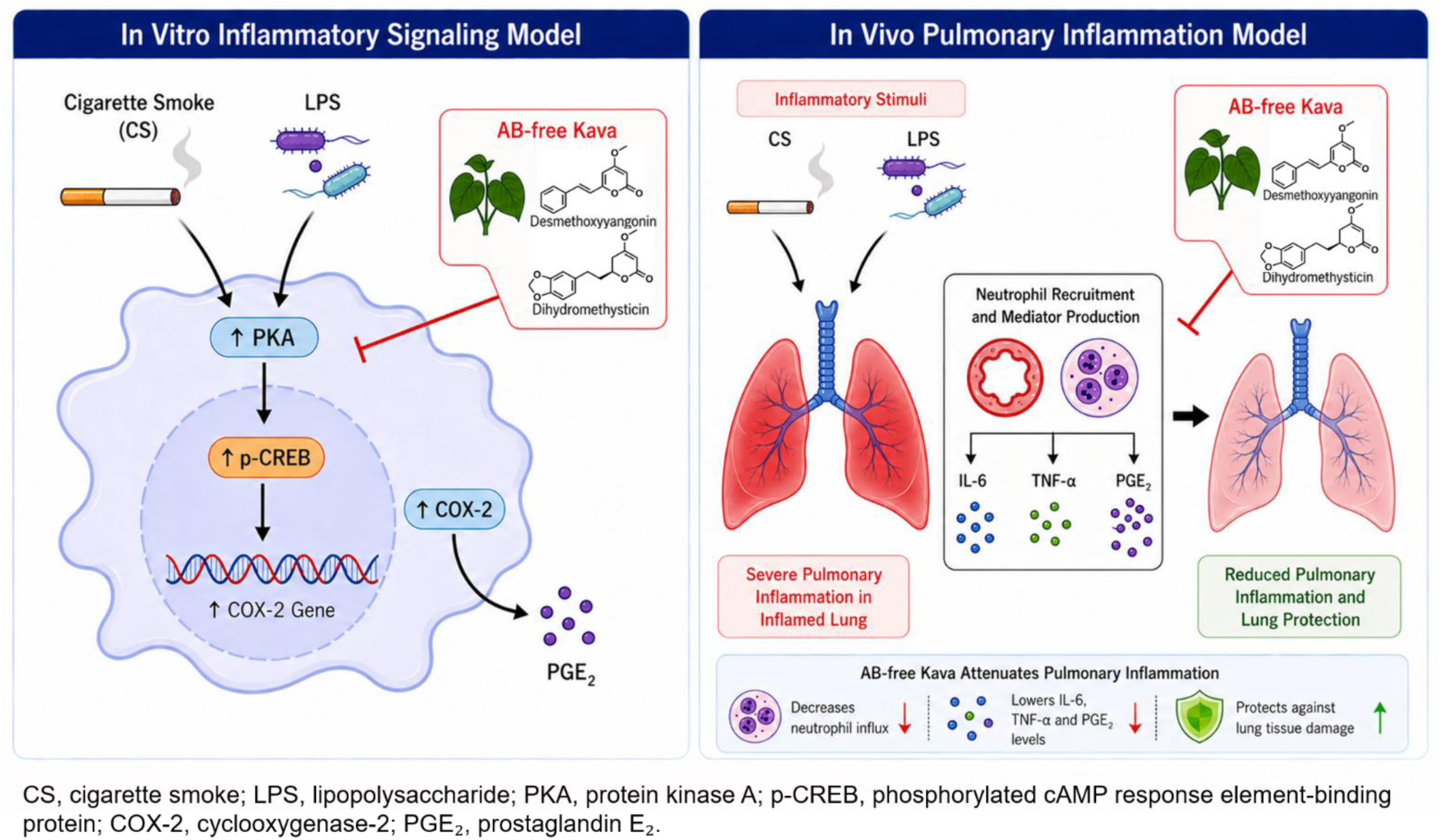

## 1. Introduction

Cigarette smoke is a major contributor to lung inflammation and the development of numerous pulmonary diseases, including chronic bronchitis, chronic obstructive pulmonary disease (COPD), lung fibrosis and lung cancer (1–4). Cigarette smoke-induced chronic inflammation is characterized by recruitment of immune cells to the airways, increased release of pro-inflammatory cytokines and chemokines, and sustained activation of transcriptional programs that amplify tissue damage and remodeling (5–7). In both clinical samples and preclinical models, cigarette smoke is closely associated with neutrophilic airway inflammation and increased inflammatory mediators, such as IL-6 and TNF-α, together with the activation of COX-2 related inflammatory pathways (8–12). Because these persistent inflammatory processes can engage in reciprocal interactions with oxidative stress and immune dysregulation, chronic lung inflammation is now viewed not simply as a downstream consequence of cigarette smoke exposure, but also as an active driver that initiates, sustains and amplifies lung disease progression (13–16).

Despite the wide availability of various anti-inflammatory medications, therapeutic control of cigarette smoke-induced lung inflammation and associated diseases remains less than optimal (17, 18). Current interventions such as NSAIDs typically show limited efficacy and can be constrained by safety concerns for long-term use (19, 20). Moreover, management of chronic inflammatory conditions requires interventions that are not only effective and safe, but also practical for sustained adherence, which highlights the appeal of diet-based strategies (“Food-Is-Medicine”) (21–23). These underscore the need for novel anti-inflammatory agents with distinct mechanisms, such as those derived from food, that can attenuate cigarette smoke-triggered inflammatory programs and ultimately reduce lung tissue injury and disease progression.

Natural products have historically provided invaluable chemical scaffolds for therapeutics in inflammation and many other diseases (23–28). Kava (*Piper methysticum*) is of particular interest because it contains a family of unique lactones, named kavalactones (Figure 1), which are structurally distinct from conventional anti-inflammatory chemotypes (29–31). Kava has revealed diverse biological activities and a number of its ingredients demonstrated anti-inflammatory potentials in cellular (32, 33) and pre-clinical animal models (34–39) while the potential of kava against cigarette smoke-induced lung inflammation was never evaluated until our recent study (40). The chemical composition of different kava products can vary substantially due to different raw materials and extraction procedures, which may influence both its efficacy and safety (30, 41). Emerging evidence suggests that kava preparations enriching flavokavains A and B (Figure 1) have increased potential of hepatotoxicity (42–48). To address these limitations, a kava formula was developed, predominantly consisting of six major kavalactones with flavokavains A and B removed, and named AB-free kava (49). AB-free kava effectively mitigated cigarette smoke-induced lung inflammation *in vivo*, reflected by reducing bronchoalveolar lavage (BAL) neutrophil infiltration and lowering pro-inflammatory TNF-α and IL-6 (40). The bioactive constituents, associated structure-activity relationship (SAR) and mechanisms remain to be defined, which is essential for clinical translation and rational optimization.

**Figure 1.**
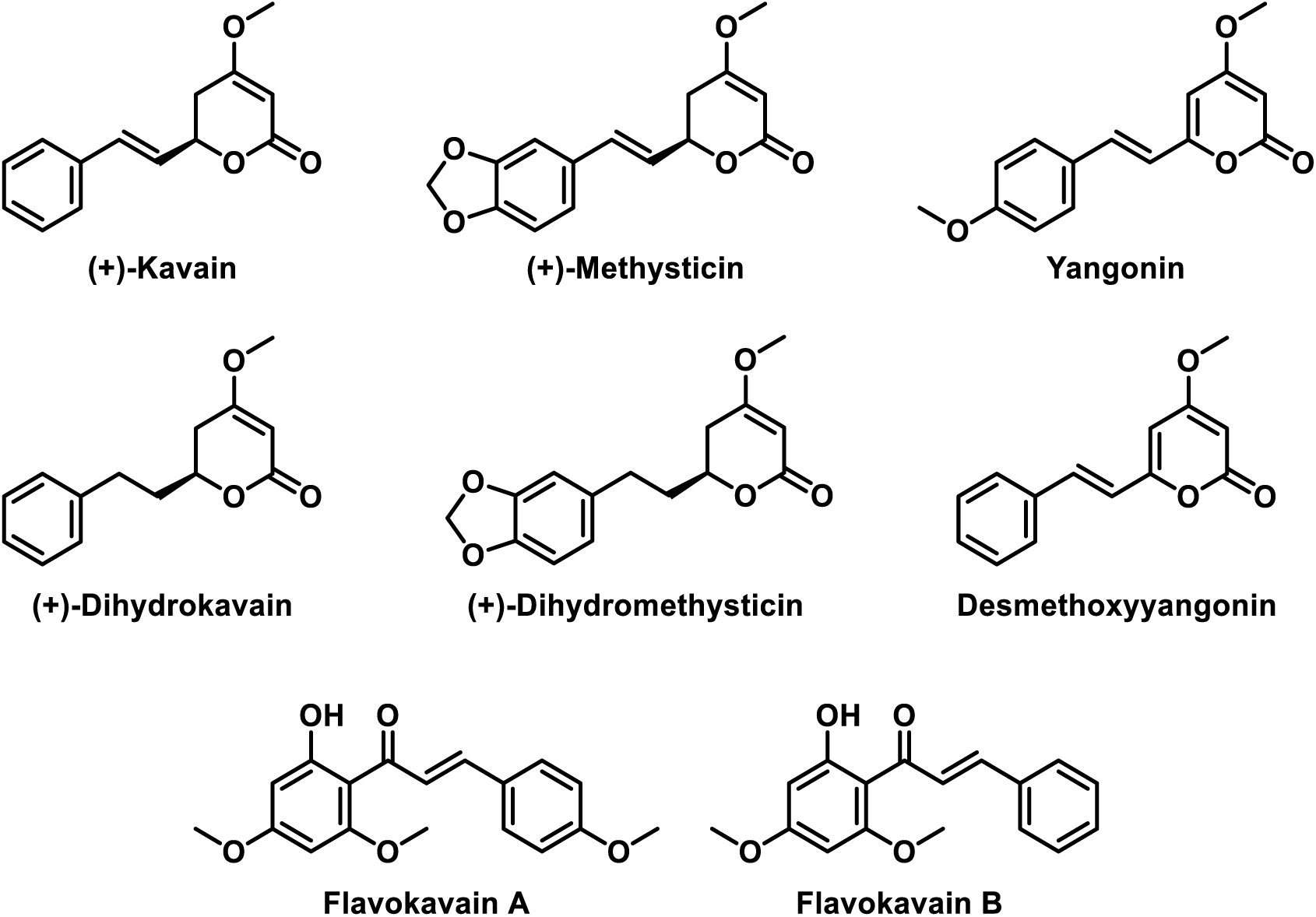
Chemical structures of six major kavalactones and flavokavains A and B in kava.

One mechanistically informative axis in inflammatory lung injury involves the induction of COX-2 and the elevation of downstream lipid-based signaling molecules (prostanoids), such as prostaglandin E₂ (PGE₂) (50, 51). COX-2 is upregulated in response to inflammatory stimuli in immune and epithelial cells and catalyzes the conversion of arachidonic acid to intermediate prostaglandin H₂, which is further processed into PGE₂ and related mediators (52–54). PGE₂ participates in immune modulation and influences the magnitude and composition of inflammatory responses by regulating phagocyte function and cytokine production (54, 55). Upstream of COX-2, multiple stimulus-responsive pathways converge to upregulate COX-2 mRNA via activating transcription factors, including nuclear factor kappa B (NF-κB), activator protein 1 (AP-1) and cAMP response element–binding protein (CREB) (56–59). Robust COX-2 induction and PGE_2_ production in macrophage systems, thus, provides a practical functional endpoint that integrates upstream transcriptional regulation and downstream mediator production (60).

In this study, we first used a lipopolysaccharide (LPS)-stimulated macrophage model to evaluate the effects of AB-free kava on suppressing LPS-induced PGE₂ production, identify the responsible kavalactones, and characterize their mode of suppression. While LPS does not mimic the full complexity of cigarette smoke exposure, it provides a robust and reproducible inflammatory stimulus for defining macrophage-centered anti-inflammatory activity. Upon identifying the *in vitro* positive lead (desmethoxyyangonin, DMY) and one negative control of a similar structure (dihydrokavain, DHK) as chemical probes, we elucidated the responsible mechanisms being protein kinase A (PKA)-medicated CREB/COX-2 signaling independent of NF-κB and AP-1. These discoveries were validated in a cigarette smoke condensate-stimulated inflammatory macrophage model and a cigarette smoke-induced lung inflammation mouse model. While the *in vitro* and *in vivo* mechanisms are highly consistent, DMY revealed moderate *in vivo* anti-inflammatory efficacy because of its low bioavailability while dihydromethysticin (DHM) revealed the best *in vivo* anti-inflammatory efficacy because of its high bioavailability despite its moderate *in vitro* potency. DHK as expected revealed minimal *in vivo* anti-inflammatory efficacy. Lastly, DHM suppressed LPS-induced lung inflammation *in vivo*. These data collectively identify DMY and DHM in AB-free kava as structurally unique and mechanistically novel anti-inflammatory therapeutic candidates that suppress LPS- and cigarette smoke-induced lung inflammation through the PKA/CREB/COX-2 axis.

## 2. Materials and Methods

### 2.1. Caution Statement for Cigarette, Cigarette Smoke and Cigarette Smoke Condensate

Cigarette, cigarette smoke (CS) and cigarette smoke condensate (CSC) are Class IA human carcinogens and should be handled carefully in well-ventilated fume hoods with appropriate protective clothing.

### 2.2. Chemicals and Reagents

AIN-93M powdered diet was purchased from Harlan Teklad (Cambridgeshire, U.K.). AB-free kava and its six major kavalactones were prepared following the procedures described in our previous study (59). For cell-based evaluations, they were dissolved in DMSO and stored at -80 °C in aliquots; each aliquot was used with no more than 2 freeze-and-thaw cycles. Diet supplemented with AB-free kava and kavalactones was prepared as previously described, stored at 4 °C, with the composition characterized every two weeks via HPLC (60). 1R6F research cigarettes were purchased from the Tobacco and Health Research Institute, University of Kentucky (Lexington, KY, USA). Cigarette smoke condensate (CSC) prepared from 3R4F reference cigarettes was purchased from Murty Pharmaceuticals (Kentucky, USA) at a stock concentration of 40 mg/mL. Lipopolysaccharide (LPS) from *Escherichia coli* O111:B4 was obtained from Sigma-Aldrich (St. Louis, MO, USA). H-89 (N-[2-((p-bromocinnamyl) amino) ethyl]-5-isoquinolinesulfonamide) was purchased from Calbiochem (San Diego, CA, USA). ELISA kits for IL-6 (M6000B) and TNF-α (MTA00B) were purchased from R&D Systems (Minneapolis, MN, USA). LC–MS grade methanol, water, acetonitrile, ethyl acetate and formic acid were purchased from Fisher Scientific (Waltham, MA, USA). Bovine serum albumin (BSA) standard was purchased from Thermo Fisher Scientific (Waltham, MA, USA). Additional reagent and antibody information was provided in Table S1.

### 2.3. Cell Culture

RAW 264.7 murine macrophages (RAW264.7) were purchased from the American Type Culture Collection (ATCC; Manassas, VA, USA) and maintained in Dulbecco’s Modified Eagle’s Medium (DMEM) supplemented with 10% fetal bovine serum (FBS) and 1% penicillin–streptomycin. Cells were cultured at 37 °C in a humidified incubator with 5% CO₂. Cells were seeded 24 h prior to treatment. After attachment, the culture medium was replaced with fresh DMEM containing 0.5% FBS for serum deprivation for 12 h. For screening experiments, cells were seeded at 5.0 × 10⁴ cells/well in 96-well plates. For Western Blotting experiments, cells were seeded at 1.0 × 10⁶ cells/well in 6-well plates. Cells were used at 80–90% confluency and were not passaged more than 10 times.

### 2.4. *In Vitro* Treatments

#### 2.4.1. LPS Stimulation of PGE_2_ production and Kavalactone Treatment

AB-free kava and individual kavalactones were prepared as stock solutions in DMSO and diluted with culture medium immediately before use. The final DMSO concentration was kept constant across all treatment groups and did not exceed 0.1% (v/v). Unless otherwise indicated, RAW 264.7 macrophages were pretreated with vehicle, AB-free kava, or individual kavalactones for 4 h, followed by co-treatment with LPS for 8 h.

For the AB-free kava dose-response study, RAW 264.7 macrophages were pretreated with AB-free kava at 1.25, 5, or 20 μg/mL for 4 h and then stimulated with increasing concentrations of LPS for 8 h. AB-free kava concentrations were based on previous clinical and preclinical studies from our group, together with preliminary optimization, to cover low, intermediate, and high non-cytotoxic exposure levels (40, 61). Culture supernatants were collected for PGE₂ quantification by LC– MS/MS. Dose-response curves were fitted by nonlinear regression, and the percentage inhibition of PGE₂ production was calculated relative to the corresponding LPS-treated group.

For the initial kavalactone screening experiment, RAW 264.7 macrophages were treated with AB-free kava (2.5 μg/mL, roughly equivalent to ∼10 μM kavalactones as the average molecular weight of the six kavalactones is roughly ∼250 Da) or one of the six major kavalactones (10 μM), including kavain (K), dihydrokavain (DHK), methysticin (M), dihydromethysticin (DHM), yangonin (Y), and desmethoxyyangonin (DMY). Cells were then stimulated with increasing concentrations of LPS for 8 h. Culture supernatants were collected for PGE₂ quantification by LC– MS/MS, and the percentage inhibition of PGE₂ production was calculated relative to the corresponding LPS-treated group.

For the kavalactone concentration-response study, RAW 264.7 macrophages were treated with increasing concentrations of each major kavalactone in the presence of LPS at 1 μg/mL for 8 h. Culture supernatants were collected and analyzed for PGE₂ levels by LC–MS/MS.

To compare the effects of DHK and DMY on LPS-induced cytokine production, RAW 264.7 macrophages were treated with DHK or DMY at 5 μM in the presence of LPS at 1 μg/mL for 24 h. Culture supernatants were collected and analyzed for IL-6 and TNF-α levels by ELISA.

#### 2.4.2. LPS-Induced PKA–CREB/COX-2 Signaling and PKA Nuclear Translocation

To characterize the time-dependent activation of inflammatory signaling pathways, RAW 264.7 macrophages were stimulated with LPS at 1 μg/mL, a concentration selected based on the LPS dose–response optimization experiment because it produced a robust PGE₂ response suitable for downstream signaling analysis, and harvested at the indicated time points. Time-matched unstimulated cells were included as controls. Cell lysates were collected for Western Blotting analysis of phosphorylated CREB, phosphorylated p65, phosphorylated c-Jun, COX-2, and related signaling proteins.

To compare the effects of individual kavalactones on LPS-induced signaling activation, cells were pretreated with DHK, DHM, or DMY at 5 μM for 4 h, followed by co-treatment with LPS at 1 μg/mL for 0.25 or 8 h. Cell lysates were collected for Western Blotting analysis, and culture supernatants were collected for PGE₂ quantification by LC–MS/MS.

For DMY concentration-response analysis, RAW 264.7 macrophages were pretreated with DMY at 1.66, 5, or 15 μM for 4 h, followed by co-treatment with LPS at 1 μg/mL for 0.25 or 8 h. Cell lysates were collected for Western Blotting analysis, and culture supernatants were collected for PGE₂ quantification by LC–MS/MS.

To assess the involvement of PKA signaling, cells were pretreated with the PKA inhibitor H89 at 2.5 or 10 μM for 1 h, followed by co-treatment with LPS at 1 μg/mL for 0.25 or 8 h. DMY at 5 μM was included as a reference treatment. Cell lysates were collected for Western Blotting analysis of CREB phosphorylation and COX-2 expression, and culture supernatants were collected for PGE₂ quantification by LC–MS/MS.

To evaluate PKA-Cα nuclear translocation, RAW 264.7 macrophages were pretreated with DHK or DMY at 10 μM for 4 h, the concentration selected based on the kavalactone screening experiment in which DMY showed clear inhibitory activity while DHK showed minimal activity and served as a structurally related negative control, followed by co-treatment with LPS at 1 μg/mL for 30 min. After treatment, cells were processed either for subcellular fractionation or immunofluorescence analysis. For subcellular fractionation, cytoplasmic and nuclear fractions were isolated and analyzed by Western blotting for PKA-Cα. α-Tubulin and Histone H3 were used as cytoplasmic and nuclear fraction markers, respectively. For immunofluorescence analysis, cells were fixed after the same treatment, stained with an antibody against PKA-Cα, and counterstained with DAPI to visualize nuclei.

#### 2.4.3. CSC Stimulation and Kavalactone Treatment

To model cigarette smoke–associated inflammatory stimulation *in vitro*, RAW 264.7 macrophages were exposed to cigarette smoke condensate (CSC) at the indicated concentrations and durations. For CSC time-course analysis, cells were treated with CSC at 20 μg/mL and harvested at the indicated time points for Western Blotting analysis of CREB phosphorylation. Time-matched unstimulated cells were included as controls. To evaluate CSC-induced COX-2 expression, RAW 264.7 macrophages were treated with CSC at 5 or 20 μg/mL and harvested at 24 or 48 h for Western Blotting analysis.

For kavalactone intervention experiments, RAW 264.7 macrophages were pretreated with vehicle, DHK, or DMY at 10 μM for 4 h, followed by co-treatment with CSC at 20 μg/mL. Cell lysates were collected at 0.25 h for p-CREB analysis and at 24 h for COX-2 analysis by Western blotting. Under the same CSC and kavalactone treatment conditions, culture supernatants were collected after 24 h for PGE₂ quantification by LC–MS/MS and for IL-6 and TNF-α measurements by ELISA.

### 2.5. PGE₂ Quantification by LC–MS/MS

Cell culture medium supernatants and BAL supernatants were collected after stimulation/exposure. For each sample, cell culture medium or BAL supernatant (75 μL) was spiked with 200 pg of PGE₂-d₄ (10 μL of 20 pg/μL) as an internal standard prior to extraction. Samples were acidified to approximately pH 3–4 with formic acid (∼200 μL of 2% formic acid) and extracted with 1 mL of ethyl acetate. The organic extracts were dried via SpeedVac and reconstituted in 30 μL of acetonitrile/water (50/50 v/v) containing 0.1% formic acid. PGE₂ was quantified by targeted UPLC–MS/MS using a Thermo Fisher Vanquish UPLC coupled to a Thermo Fisher Altis Plus triple quadrupole mass spectrometer. Chromatographic separation was performed on an Atlantis dC18 column (50 × 2.1 mm, 3 μm; Waters) using a 7.5 min gradient at a flow rate of 0.250 mL/min, with an injection volume of 15 μL. Mobile phase A was 0.1% formic acid in water, and mobile phase B was 0.1% formic acid in acetonitrile. The column temperature was maintained at 30 °C, and the autosampler was maintained at 5 °C. The gradient started at 35% B for 0.5 min, increased to 95% B over 2 min, held at 95% B for 1.5 min, and then returned to 35% B for the remainder of the run for column equilibration. The mass spectrometer was operated in negative ionization mode with HESI source settings as follows: spray voltage, 3345 V; sheath gas, 50; auxiliary gas, 10; sweep gas, 0; ion transfer tube temperature, 325 °C; vaporizer temperature, 350 °C; and CID gas pressure, 1.5 mTorr. Detection was performed in selected reaction monitoring (SRM) mode using multiple transitions. For PGE₂, the quantification transition was m/z 351.160 → 271.259 (CE = 16), and a qualifying transition of m/z 351.160 → 315.229 was monitored for identity confirmation. For PGE₂-d₄, the quantification transition was m/z 355.280 → 275.279 (CE = 16), and a qualifying transition of m/z 355.280 → 319.219 was monitored. At least three technical replicates were performed with the mean and standard deviation reported. The calibration curve, limit of detection (LOD), and limit of quantification (LOQ) were provided in the Supporting Information (Figure S1).

### 2.6. Cytokine Measurements by ELISA

Cell culture medium supernatants and BAL supernatants were clarified by centrifugation at 300 × g for 5 min and stored at −80 °C until analysis. IL-6 and TNF-α levels were quantified using ELISA kits from R&D Systems (IL-6: M6000B; TNF-α: MTA00B) according to the manufacturers’ instructions. Concentrations were calculated from standard curves generated in parallel. At least three technical replicates were performed with the mean and standard deviation reported.

### 2.7. Western Blotting of Cell and Lung Tissue Samples

For cell-based experiments, cell pellets (approximately 1 × 10⁶ cells per sample) were collected after treatment and lysed in 50 μL RIPA buffer supplemented with protease and phosphatase inhibitors. Lysates were clarified by centrifugation at 13,000 × g for 15 min at 4 °C, and the supernatants were collected. For lung tissue experiments, lung tissue (20 mg) was homogenized in 250 μL RIPA buffer supplemented with protease and phosphatase inhibitors, and the supernatants were collected after centrifugation at 13,000 × g for 15 min at 4 °C. Protein concentrations were determined using a BCA assay. Equal amounts of protein were subjected to SDS–PAGE (30 μg for cell lysates and 50 μg for lung tissue lysates) using 4–12% Bis–Tris gels, followed by transfer to PVDF membranes. Membranes were blocked with 5% non-fat milk in TBST for 1 h at room temperature (or with 5% milk in TBST as indicated) and incubated with primary antibodies (Table S1) overnight at 4 °C. Membranes were then incubated with the corresponding HRP-conjugated secondary antibodies for 1.5–2 h at room temperature. Protein bands were detected using enhanced ECL reagent (Thermo Scientific) and visualized using a Bio-Rad ChemiDoc imaging system. At least three replicates were performed for each protein target and representative blots are shown.

### 2.8. Nuclear and Cytoplasmic Fractionation for Assessment of PKA Nuclear Translocation

To assess PKA-Cα nuclear translocation, treated RAW 264.7 cells were fractionated into cytoplasmic and nuclear extracts using NE-PER Nuclear and Cytoplasmic Extraction Reagents (Thermo Scientific, Waltham, MA, USA) according to the manufacturer’s instructions. Briefly, cell pellets were harvested after treatment and lysed in cytoplasmic extraction reagent supplemented with protease and phosphatase inhibitors. After centrifugation, the resulting supernatants were collected as the cytoplasmic fraction, whereas the pellets containing nuclei were further extracted with nuclear extraction reagent to obtain nuclear proteins. Protein concentrations were determined by BCA assay, and equal amounts of protein from each fraction were analyzed by SDS–PAGE and Western blotting as described above. PKA-Cα was detected to assess its subcellular distribution, with α-tubulin and histone H3 serving as cytoplasmic and nuclear markers, respectively. At least three independent experiments were performed with representative blots shown.

### 2.9. Immunofluorescence Assessment of PKA Nuclear Translocation

For PKA nuclear translocation, treated cells were fixed with 4% paraformaldehyde for 15 min and permeabilized with 1% Triton X-100 in PBS for 10 min, followed by blocking with 5% BSA in PBS for 1 h. After blocking, cells were incubated with PKA-Cα antibody overnight at 4 °C and secondary antibody for 1 h at room temperature. Nucleus was stained with DAPI for 15 min. Cells were imaged using fluorescence microscopy (Nikon Ti2, Japan). At least three replicates were performed with representative images shown.

### 2.10. Animal Study

#### Cigarette smoke-induced lung inflammation in A/J mice

Female A/J mice (6 weeks old, 16–18 g) were purchased from The Jackson Laboratory (Bar Harbor, ME, USA) and maintained in specific pathogen-free facilities under animal welfare protocols approved by the Institutional Animal Care and Use Committees of the University of Florida (IACUC protocol #202300000018). Mice were housed with ad libitum access to water and AIN-93M powdered diet (basal diet) and allowed to acclimate for 7–10 days before study initiation. After acclimation, mice were weighed and randomized based on bodyweight into six experimental groups (n = 8/group): filtered air control, cigarette smoke (CS) only, CS + AB-free kava (1000 ppm in diet), CS + DHM (200 ppm in diet), CS + DMY (200 ppm in diet), and CS + DHK (200 ppm in diet). For dietary supplementation experiments, mice were maintained on AIN-93M powdered diet with or without the indicated supplements throughout the exposure period. Bodyweight was measured weekly, and cage-level food intake was estimated twice weekly.

CS exposure sessions were conducted using 1R6F research cigarettes via a microprocessor-controlled cigarette smoking machine (model TE-10, Teague Enterprises, Davis, CA, USA) according to the Federal Trade Commission (FTC) smoking protocol. Mice were exposed to CS for 2 h/day (9:00–11:00 a.m.), Monday through Friday, for 2 weeks (i.e., no weekend exposure). Cigarettes were smoked using a standardized procedure (35 cm³ puff volume, 1 puff/min, 2 s per puff, 8 puffs/cigarette). A mixture of mainstream and sidestream smoke was conveyed to a mixing/diluting chamber, aged for several minutes, and diluted with air to approximately 100 mg/m³ total suspended particles (TSP). During each daily CS exposure session, TSP and CO levels were monitored. TSP was determined by gravimetric sampling, and CO levels were measured using a calibrated CO analyzer (Monoxor III, Bacharach, Kensington, PA, USA). Air-control mice were handled in parallel and maintained under filtered-air exposure conditions. Mice were euthanized by CO₂ asphyxiation approximately 22 h after the final CS exposure. At necropsy, BAL, blood (for serum preparation), urine, and tissues including lung, liver, and heart were collected for analysis.

#### LPS-induced lung inflammation in C57BL/6J mice

Female C57BL/6J mice (10 – 12 weeks old, 25 – 30 g) were purchased from The Jackson Laboratory (Bar Harbor, ME, USA) and maintained in specific pathogen-free facilities under animal welfare protocols approved by the Institutional Animal Care and Use Committees of the University of Florida (IACUC protocol #202300000018). Mice were housed with ad libitum access to water and diet and allowed to acclimate for 7–10 days before study initiation. After acclimation, mice were weighed and randomized based on bodyweight into three experimental groups (n = 5/group): LPS only, LPS + DHM (40 mg per kg bodyweight), and LPS + DHK (40 mg per kg bodyweight). DHM and DHK were given to mice via oral gavage (0.2 mL corn oil) 1 h before LPS challenge with corn oil given to the control mice. LPS (0.25 μg/mouse) were administered via intrachial injection. Mice were euthanized 24 h after LPS exposure with BAL collected and analyzed.

### 2.11. Bronchoalveolar Lavage Collection

The trachea of each euthanized mouse was surgically exposed and cannulated with 27-gauge silicone tubing connected to a 23 G needle on a 1 mL syringe. Sterile PBS (1.0 mL) was instilled into the lungs via the trachea and gently withdrawn to BAL. The samples were centrifuged at 300× g for 5 min to separate cells from supernatant. Cells were analyzed by flow cytometry while the BAL supernatant was collected and stored at −80 °C until analysis.

### 2.12. Immune Cell Profiling Assays

BAL cells were collected as described above, washed with PBS, and incubated with Fc receptor blocker (anti-mouse CD16/CD32, clone 2.4G2, BD Biosciences) together with Zombie Red Fixable Viability Dye (1:500). Cells were then stained with fluorophore-conjugated antibodies against CD45, CD11b, and Ly6G for neutrophil analysis. Flow cytometry was performed using a Cytek Aurora cytometer (Cytek Biosciences, Fremont, CA, USA), and the resulting data were analyzed using FlowJo software (BD Biosciences). Neutrophils were identified from live CD45⁺ leukocytes based on CD11b⁺Ly6G⁺ staining. Flow cytometric analysis was carried out according to our previously reported protocols(40).

### 2.13. TNF-α and IL-6 Measurements in BAL

TNF-α and IL-6 levels in BAL supernatant samples were quantified using ELISA kits from R&D Systems (TNF-α: MTA00B; IL-6: M6000B-1) according to the manufacturer’s recommended protocols. Samples were assayed in duplicate. Concentrations were calculated from standard curves generated in parallel.

### 2.14. Statistical Analysis

Data were reported as the mean ± standard deviation (SD). One-way ANOVA was used to compare the means among multiple treatment groups. For comparison between two groups, a two-tailed Student t test was employed. A p value of < 0.05 was considered statistically significant. All analyses were conducted using GraphPad Prism 10 (GraphPad Software, Inc.).

## 3. Results

### 3.1. AB-Free Kava Suppresses LPS-Induced PGE₂ Production in Macrophages with Desmethoxyyangonin (DMY) as One Positive Lead and Dihydrokavain (DHK) as One Negative Control

The macrophage-based PGE_2_ platform was established to evaluate AB-free kava and to identify kavalactone(s) capable of suppressing COX-2-linked inflammatory mediator output. To optimize assay conditions for screening, the dose-response LPS-induced PGE₂ production in RAW 264.7 macrophages was first established (Figure S1A). Time-course analysis at the selected LPS concentration (1 μg/mL) showed that PGE₂ accumulation peaked at approximately 8 h (Figure S1B), establishing the time window suitable for screening kavalactone-mediated suppression. This time point is consistent with previous studies using an 8 h LPS stimulation period to evaluate PGE₂ production in RAW 264.7 macrophages (62). Under these optimized conditions, AB-free kava markedly reduced LPS-induced PGE₂ accumulation across the tested LPS concentrations (Figure 2A), indicating that AB-free kava contains bioactive constituents capable of attenuating inflammatory mediator production during LPS-driven macrophage activation. The selected LPS concentrations covered a broad range, with the lowest doses (0.00001 and 0.0001 µg/mL) falling within or close to the clinically relevant endotoxin range, whereas the higher doses were included as supraphysiologic concentrations to elicit robust inflammatory responses *in vitro* (63). Specifically, there was a greater suppression at lower LPS concentrations with the low dose of AB-Free kava (Figure 2B), which is more likely to represent physiological LPS exposure and human kava exposure. To identify the constituents responsible for this activity, six major kavalactones (kavain – K, dihydrokavain – DHK, methysticin – M, dihydromethysticin – DHM, yangonin – Y, and desmethoxyyangonin – DMY) were evaluated at a fixed concentration (10 µM) with LPS dose-escalation. Distinct inhibitory profiles were observed with a clear structure–activity relationship (SAR). Specifically, DMY exhibited the strongest suppression of LPS-induced PGE₂ followed by Y. M and DHM revealed weaker but similar inhibitory effects while K and DHK showed minimal activity under the same conditions (Figure 2C). Most of these kavalactones exhibited significant suppression of LPS-induced PGE_2_ production at the lowest LPS concentration, indicating their anti-inflammatory potential likely being physiologically relevant (Figure 2D). Lastly, the dose-response effects of six kavalactones were analyzed at a fixed LPS concentration (1 µg/mL), which again identified DMY as the most effective inhibitor among the tested kavalactones (Figure 2E). The effects of DHK and DMY on LPS-induced TNF-α and IL-6 production were also assessed (Figure 2F), further supporting their differential anti-inflammatory potential. Given that DMY and DHK share high structural similarity and distinct inhibitory activity against LPS-induced inflammatory mediator production, they were selected as chemical probes for subsequent mechanistic interrogation. DHM was also selected as a lead compound because, despite its intermediate *in vitro* activity (Figures 2C and 2E), it exhibited higher *in vivo* bioavailability than DMY, raising the possibility that the most potent *in vitro* candidate may not necessarily represent the best *in vivo* candidate. (64).

**Figure 2.**
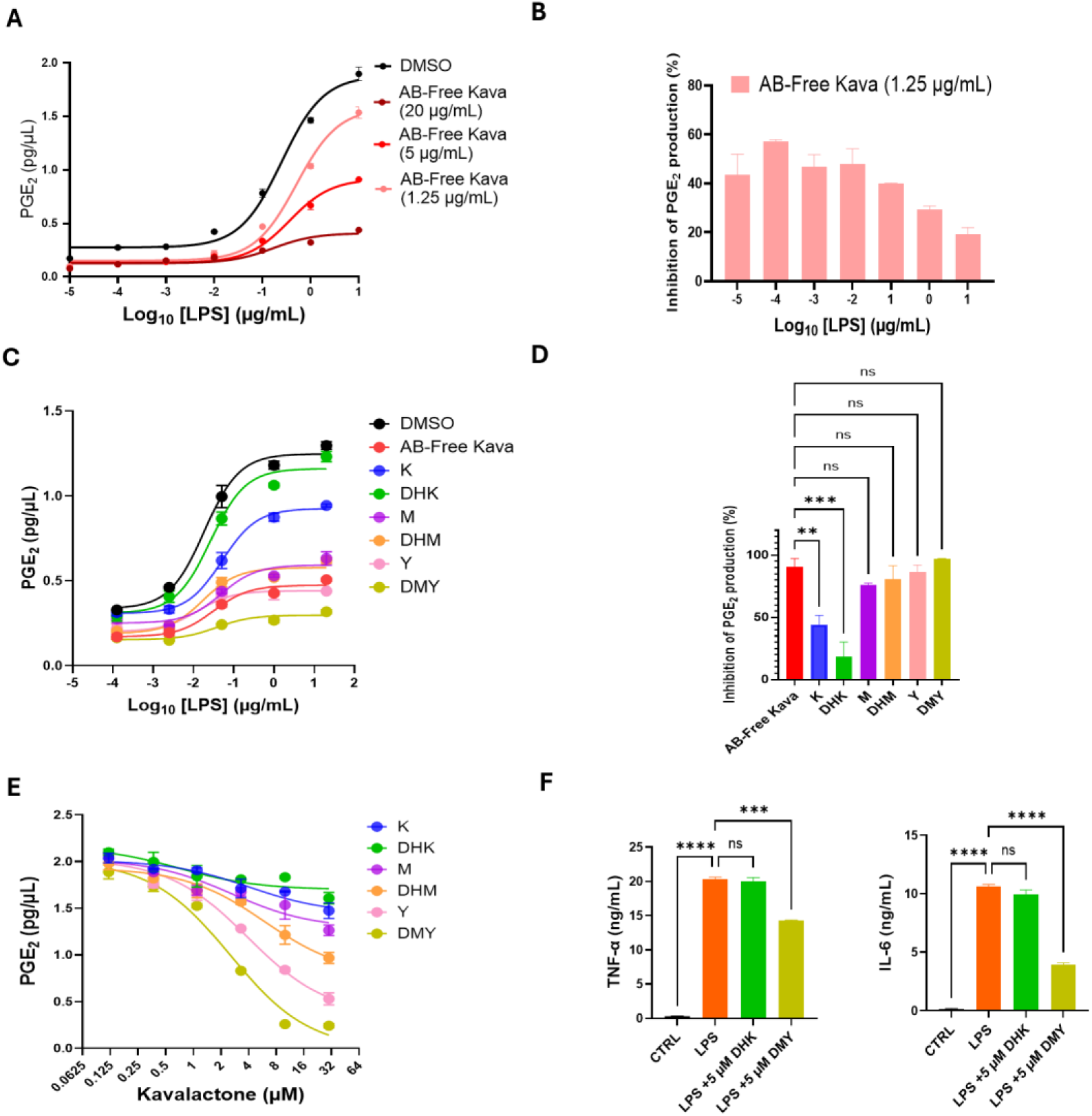
AB-free kava and its major kavalactones suppress LPS-induced PGE₂ production in RAW 264.7 macrophages with DMY as a lead inhibitor and DHK as a negative probe. (A) Dose-dependent effect of AB-free kava on PGE₂ production across increasing LPS concentrations; dose–response curves were fit by nonlinear regression. (B) The percentage of PGE_2_ inhibition by AB-free kava (1.25 μg/mL) relative to the corresponding LPS treatment. (C) Comparative effects of AB-free kava (2.5 μg/mL) and six major kavalactones (10 μM) on LPS-induced PGE₂ production under LPS dose-escalation conditions. (D) The percentage of PGE_2_ inhibition by different kavalactones relative to the corresponding LPS treatment (10^-4^ μg/mL). (E) Dose-response analysis of the six major kavalactones against LPS (1 μg/mL)-induced PGE₂ production. (F) Comparative effects of DHK and DMY on LPS (1 μg/mL)-induced IL-6 and TNF-α release. Data are presented as mean ± SD (n = 3); in some panels, error bars are not visible because of minimal variation among replicates. Statistical analysis was performed using one-way ANOVA with a Dunnett’s multiple comparisons test; * p < 0.05; ** p < 0.01; *** p < 0.001; **** p < 0.0001; ns, not significant.

### 3.2. Time-Course Analysis Defines a Temporal Window Associated with LPS-Induced COX-2 Induction and Potential Upstream Signaling Pathways

Given the established connection between COX-2 induction and PGE₂ production, signaling kinetics were profiled after LPS stimulation to define appropriate time points for mechanistic studies and to identify potential upstream regulatory pathways. Because COX-2 is primarily induced at the transcriptional level upon inflammatory stimulation, the activation of three transcription factors (NF-κB, AP-1 and CREB) as potentially responsible upstream signaling were evaluated by monitoring the phosphorylation of p65, c-Jun and CREB respectively (65). Phosphorylation of p65, c-Jun, and CREB were observed within 0.25 h upon LPS treatment and appeared to sustain (Figure 3A). As expected, COX-2 protein induction was delayed, becoming evident at approximately 4 h and reaching maximal levels by 8 h (Figure 3A, longer timepoint data not shown), supporting a temporal sequence in which upstream signaling precedes COX-2 induction. Based on these data, 0.25 h was selected as the timepoint to capture early signaling events, whereas 8 h was used as a downstream timepoint to quantify COX-2 induction for subsequent experiments.

**Figure 3.**
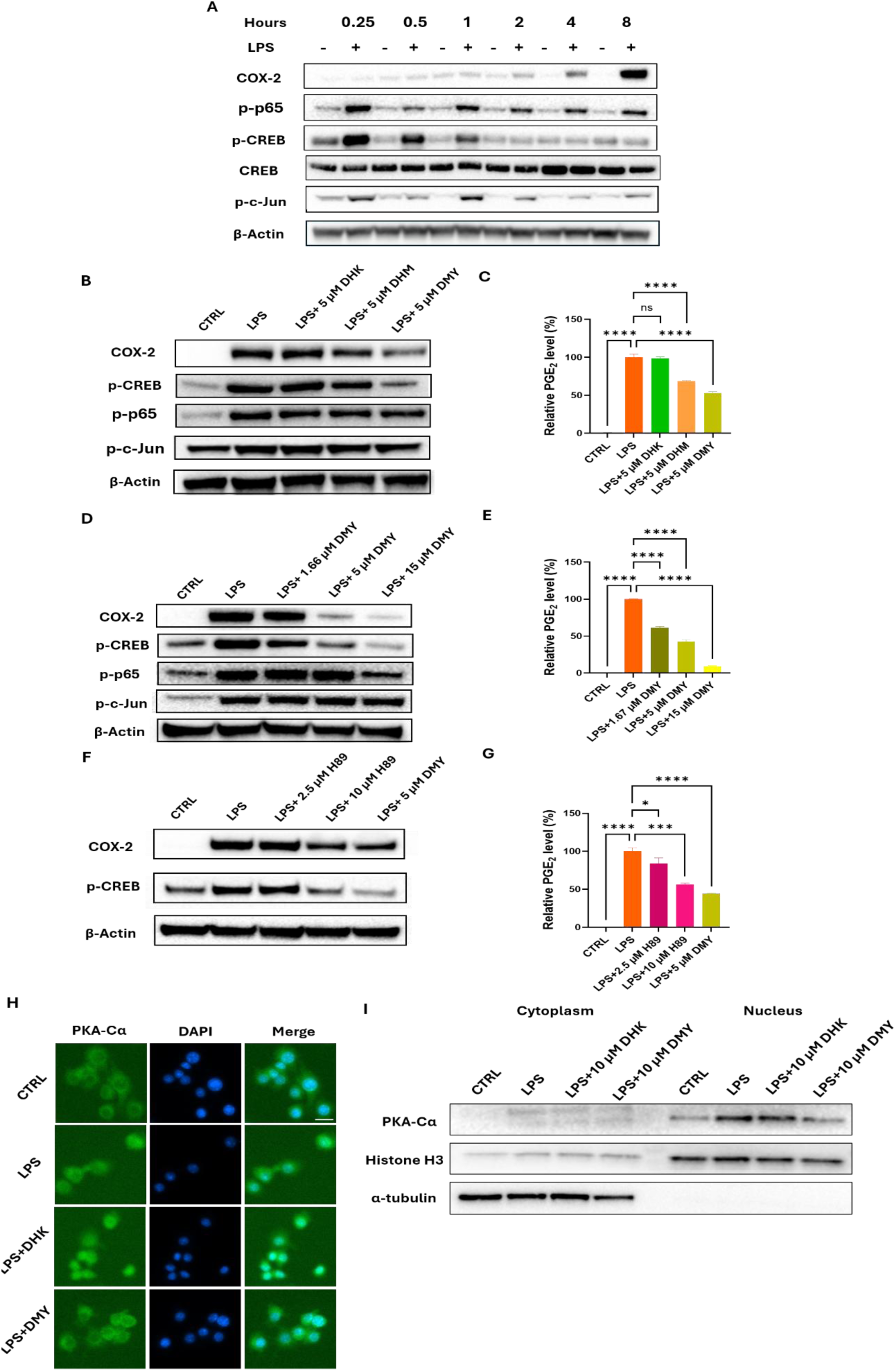
Kavalactones modulate the PKA–CREB/COX-2 axis in LPS-stimulated RAW 264.7 macrophages. (A) Time-course analysis of LPS-induced PKA–CREB/COX-2 signaling activation in RAW 264.7 macrophages. The “−” lanes indicate time-matched unstimulated controls. (B) Comparative effects of DHK, DHM, and DMY on LPS-induced inflammatory signaling, including CREB, NF-κB p65, and c-Jun phosphorylation, as well as COX-2 expression. (C) PGE₂ levels in culture supernatants from the conditions in (B) were quantified by LC–MS/MS and expressed as relative levels. (D) Western Blotting analysis showing the concentration-dependent effect of DMY on LPS-induced CREB phosphorylation and COX-2 expression. (E) PGE₂ levels in culture supernatants from the conditions in (D) were quantified by LC–MS/MS and expressed as relative levels. (F) Western Blotting analysis showing the effect of the PKA inhibitor H89 on LPS-induced CREB phosphorylation and COX-2 expression, with DMY included as a reference treatment. (G) PGE₂ levels in culture supernatants from the conditions in (F) were quantified by LC–MS/MS and expressed as relative levels. (H) Representative immunofluorescence images showing LPS-induced nuclear translocation of PKA-Cα and its attenuation by kavalactone treatment. PKA-Cα is shown in green, and nuclei are counterstained with DAPI in blue. Representative immunofluorescence images from NC, LPS, LPS + DHK, and LPS + DMY groups are shown. Scale bar 10 μm. (I) Western Blotting analysis of PKA-Cα distribution in cytoplasmic and nuclear fractions, showing LPS-induced nuclear accumulation of PKA-Cα and its modulation by kavalactone treatment. α-Tubulin and Histone H3 were used as cytoplasmic and nuclear markers, respectively. Data are presented as mean ± SD. Statistical analysis was performed using one-way ANOVA with Dunnett’s multiple comparisons test; *p < 0.05; **p < 0.01; ***p < 0.001; ****p < 0.0001; ns, not significant.

### 3.3. DMY, DHM, and DHK differentially suppress CREB/COX-2/PGE₂ signaling but not p65 or c-Jun activation in LPS-stimulated macrophages

To explore the responsible upstream transcriptional programs, representative kavalactones (DHK, DHM, and DMY) were compared under matched LPS stimulation conditions, and signaling readouts linked to COX-2 regulation were assessed. DMY, DHM and DHK treatments showed distinct and consistent effects on LPS-induced p-CREB, COX-2 and PGE₂ (DMY > DHM > DHK), whereas none of them have significant effects on NF-κB (p-p65) and AP-1 (p-c-Jun) signaling (Figure 3B, C). These results support CREB phosphorylation as a potential up-stream signaling associated with kavalactone-mediated suppression of COX-2-linked inflammatory outputs.

DMY further demonstrated a dose-dependent suppression of COX-2 induction, p-CREB and PGE_2_ production while it had no effects on p-p65 nor p-c-Jun at the lower concentrations (1.66 and 5 µM), which already resulted in significant reductions in COX-2 and PGE_2_ (Figure 3D and 3E). The similar concentration-dependent reductions in p-CREB, COX-2 and PGE_2_ further supports a mechanism in which DMY attenuates a CREB-associated inducible transcriptional program that contributes to COX-2 expression and downstream PGE₂ production, providing mechanistic coherence with the functional PGE₂ suppression observed in the screening experiments while NF-κB (p-p65) and AB-1 (p-c-Jun) were again unlikely involved (Figure 3D).

### 3.4. Pharmacological Inhibition Supports PKA as a Major Upstream Regulator of the CREB/COX-2/PGE_2_ Axis

Given the canonical role of PKA in regulating CREB phosphorylation, pharmacological PKA inhibition was used to evaluate whether PKA lies upstream of the CREB/COX-2 program in this model (58, 66). In LPS-stimulated RAW 264.7 macrophages, H89 (a PKA inhibitor) effectively reduced CREB phosphorylation and attenuated COX-2 induction and PGE_2_ production (Figure 3F and 3G), similar as DMY, supporting PKA activation as a potential up-stream signaling for kavalactones (67). To further assess PKA activation, the subcellular localization of the PKA catalytic subunit (PKA-Cα) was examined by both immunofluorescence analysis and subcellular fractionation. Immunofluorescence imaging showed that LPS treatment promoted the redistribution of PKA-Cα from the cytoplasm to the nucleus, indicating PKA activation, and this nuclear translocation was attenuated by DMY treatment (Figure 3H). Consistently, subcellular fractionation analysis further confirmed that LPS increased PKA-Cα levels in the nuclear fraction, whereas DMY treatment partially reversed this effect (Figure 3I). Together, these results suggest that DMY suppresses LPS-induced PKA activation while DHK was not effective. Collectively, these data indicate that LPS exposure induces a PKA-dependent CREB/COX-2 axis, contributing to inducible COX-2 expression and downstream PGE₂ production while kavalactones blocks PKA activation and thus suppresses the down-stream signaling and associated inflammation.

### 3.6. Kavalactones Suppress CSC-Induced CREB/COX-2 Activation and Cytokine Production

To complement the LPS-based screening platform with a cigarette smoke-relevant stimulus, CSC was used to stimulate RAW 264.7 macrophages. Although much weaker than LPS stimulation, CSC exposure (20 μg/mL) rapidly induced CREB phosphorylation at 0.25 h (Figure 4A), followed by slower COX-2 induction at 24 h after CSC exposure (Figure 4B). DMY and DHK were therefore evaluated under CSC stimulation (20 μg/mL) using these respective time points for mechanistic assessment. DMY attenuated CSC-induced CREB phosphorylation at 0.25 h and reduced COX-2 expression at 24 h, whereas DHK showed weaker effects (Figure 4C). Consistent with its stronger suppression of COX-2, DMY markedly inhibited CSC-induced PGE₂ production, whereas DHK showed only a modest inhibitory effect (Figure 4D). DMY also more effectively reduced CSC-induced IL-6 and TNF-α release, whereas DHK produced weaker attenuations under the same conditions (Figure 4E). Together, these results suggest that CSC also drives inflammatory cytokine release and PGE_2_ production through modulating the same CREB/COX-2 signaling cascade as observed in the LPS model and in both models, kavalactones could suppress such signaling, mechanistically responsible for their anti-inflammatory potential with DMY being more potent than DHK.

**Figure 4.**
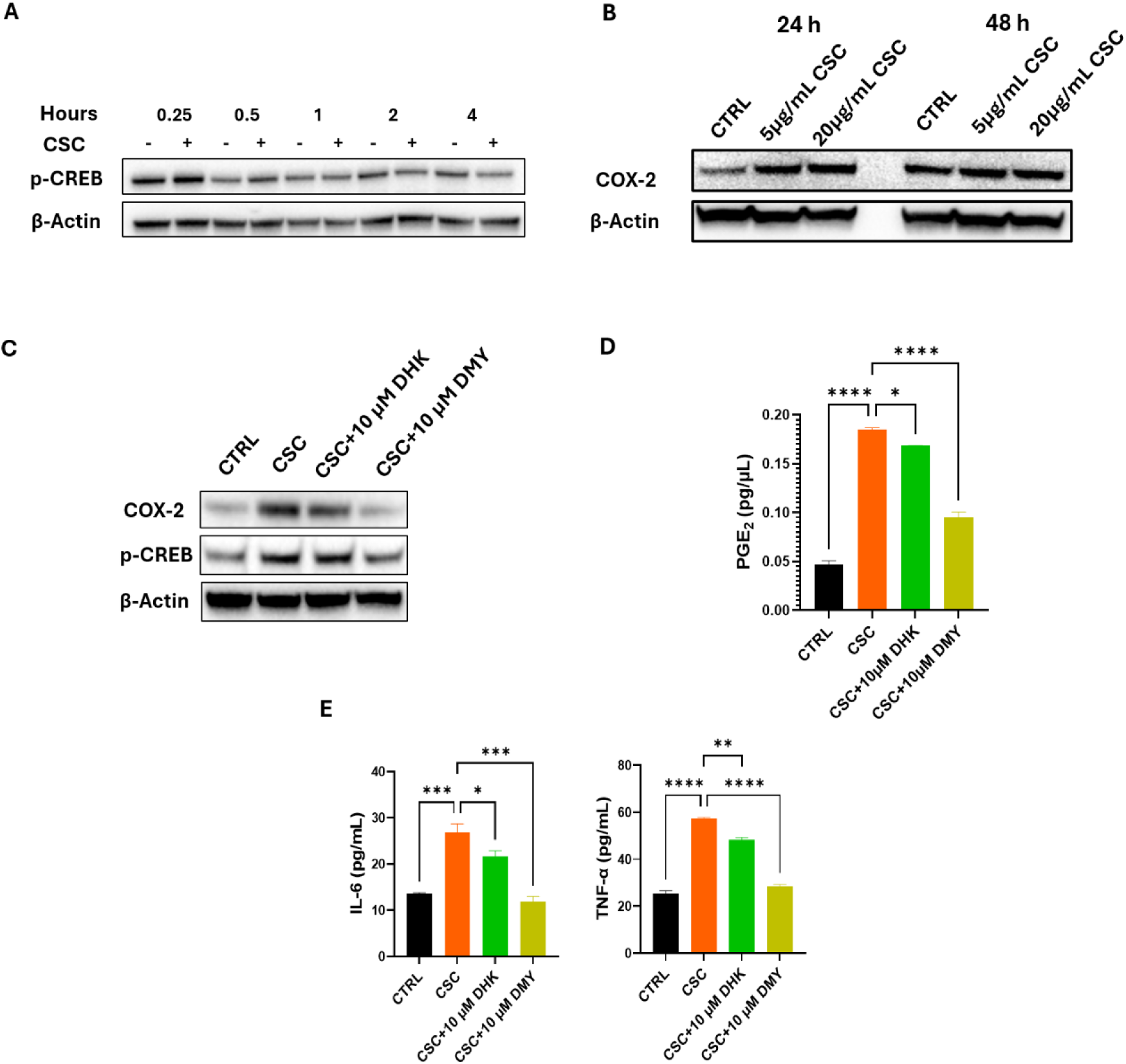
Kavalactones suppress CSC-induced PGE₂ production and cytokine release and attenuate CSC-driven activation of the CREB/COX-2 axis in RAW 264.7 macrophages. (A) Time-course analysis of CSC-induced CREB phosphorylation in RAW 264.7 macrophages. The “−” lanes indicate time-matched unstimulated controls. (B) Dose- and time-dependent induction of COX-2 expression by CSC. (C) Effects of DHK and DMY on CSC-induced COX-2 expression and CREB phosphorylation. Cell lysates were collected at 0.25 h for p-CREB analysis and at 24 h for COX-2 analysis by immunoblotting. (D) Relative PGE₂ levels in culture supernatants following CSC stimulation with or without DHK or DMY treatment, as quantified by LC–MS/MS. (E) Under the same treatment conditions, IL-6 and TNF-α levels in culture supernatants were quantified by ELISA. Data are presented as mean ± SD. Statistical analysis was performed using one-way ANOVA with Dunnett’s multiple comparisons test; *p < 0.05; **p < 0.01; ***p < 0.001; ****p < 0.0001; ns, not significant.

### 3.7. AB-Free Kava and Kavalactones Mitigate Cigarette Smoke–Induced Lung Inflammation in Mice and Modulate Lung CREB/COX-2 Signaling

Cigarette smoke exposure elicited a pronounced inflammatory phenotype *in vivo*, characterized by increased airway neutrophilia and elevated inflammatory mediators in BAL (68), which was observed in A/J mice upon a 2-week cigarette smoke exposure (Figure 5A and 5B). The effects of AB-free kava and individual kavalactones (DMY, DHK and DHM) were evaluated via dietary supplementation. DMY was evaluated *in vivo* because of its potent *in vitro* activity while DHK was evaluated as a negative control for its minimal *in vitro* activity. Our previous pharmacokinetic study, on the other hand, revealed minimal bioavailability for DMY while DHM have much better bioavailability (64). DHM therefore was evaluated here as well despite its moderate *in vitro* activity. AB-free kava was evaluated at a dose of 1 mg/g of diet while individual kavalactones were evaluated at a dose of 0.2 mg/g of diet given that each individual kavalactones accounts for 10 – 40% of the mass of AB-free kava. Consistent with our previous study, AB-free kava strongly reduced cigarette smoke-induced BAL neutrophils, IL-6, TNF-α and PGE_2_ to a level similar to mice without cigarette smoke exposure (Figure 5A and 5B). Among the three kavalactone evaluated, DHK showed no inhibitory effects on neutrophil accumulation and the levels of IL-6 and TNF-α in BAL. Although it significantly reduced the levels of PGE_2_, its efficacy appeared to be weaker than the other two kavalactones. DMY significantly reduced all inflammatory measures except IL-6 and it was not as effective as AB-free kava. DHM at the same time showed similar efficacy as AB-free kava, much more effective than DMY, likely because of the much better *in vivo* bioavailability of DHM in comparison to DMY. Consistent with the BAL inflammatory measures, immunoblot analysis of lung tissue indicated that cigarette smoke increased COX-2 expression and CREB phosphorylation, whereas AB-free kava and kavalactones reduced these smoke-associated elevations with the levels of reduction overall positively correlated with their suppression of inflammatory measures that AB-free kava and DHM showed the most significant suppression followed by DMY while DHK was not effective (Figure 5C). Collectively, these findings demonstrate that AB-free kava and kavalactones, particularly DHM and DMY, effectively suppress cigarette smoke–induced airway inflammation and inflammatory mediator production *in vivo*, accompanied by attenuation of lung CREB/COX-2 signaling.

**Figure 5.**
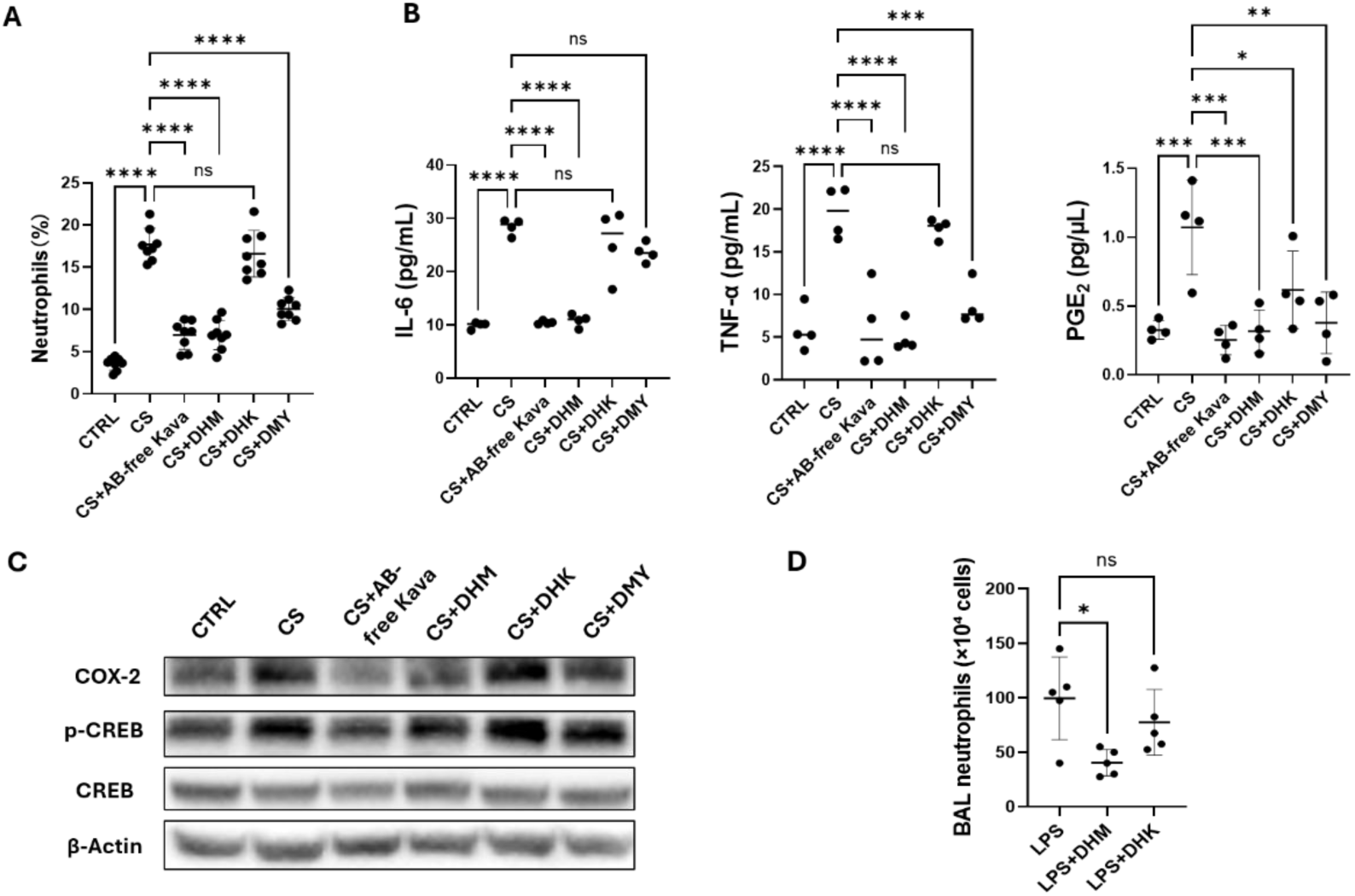
AB-free kava and kavalactones mitigate cigarette smoke– and LPS-induced lung inflammation in mice and attenuate lung CREB/COX-2 signaling. (A) Flow cytometric quantification of neutrophils in bronchoalveolar lavage (BAL) fluid from A/J mice. Data is presented as mean ± SD (n = 8 mice per group). (B) BAL concentrations of IL-6 and TNF-α were quantified by ELISA, and BAL PGE₂ levels were quantified by LC–MS/MS (n = 4 mice per group). Data are presented as mean ± SD. (C) Western Blotting analysis of lung tissue lysates for COX-2, phosphorylated CREB (p-CREB) and total CREB with β-actin as a loading control. (D) Effects of DHM and DHK on LPS-induced neutrophil accumulation in BAL from female C57BL/6J mice. DHM and DHK were administered by oral gavage at 40 mg/kg body weight 1 h before LPS challenge, and BAL neutrophils were quantified 24 h after LPS exposure. Data is presented as mean ± SD. Statistical analysis was performed using one-way ANOVA followed by Dunnett’s multiple comparisons test.; *p < 0.05; **p < 0.01; ***p < 0.001; ****p < 0.0001; ns, not significant.

### 3.8. DHM not DHK Mitigates LPS–Induced Neutrophil Accumulation in BAL in C57BL/6J mice

We lastly evaluated the effects of DHM and DHK on LPS-induced lung inflammation in female C57BL/6J mice. DHM and DHK were administered via oral gavage at 40 mg/kg bodyweight 1 h before LPS challenge. BAL neutrophils were quantified 24 h after LPS exposure. As shown in Figure 5D, DHM significantly reduced LPS-induced neutrophil accumulation in BAL, whereas DHK did not show a significant effect. These findings further support DHM as a major *in vivo* anti-inflammatory constituent of AB-free kava.

## 4. Discussion and conclusion

Cigarette smoke-induced lung inflammation is a major driver of airway injury and disease progression, yet available anti-inflammatory strategies often provide incomplete protection or are limited by safety. Kava (*Piper methysticum*) contains structurally distinct kavalactones, offering an attractive natural-product scaffold for identifying anti-inflammatory agents with potentially new modes of action. Motivated by our prior observations that AB-free kava mitigates cigarette smoke–induced lung inflammatory responses in mice, this study integrated constituent screening, mechanistic interrogation in macrophages, smoke-relevant *in vitro* stimulation, and *in vivo* validation to have identified the bioactive kavalactones (DHM and DMY) and elucidated a coherent signaling framework (PKA-mediated CREB/COX-2 axis) that links kavalactone exposure to suppression of inflammatory outputs across multiple experimental contexts.

A central mechanistic finding is that kavalactones can attenuate PKA/CREB/COX-2–linked inflammatory program associated with reduced inflammatory mediator production. In LPS-stimulated RAW 264.7 macrophages, AB-free kava robustly suppressed PGE₂ production. Screening of six major kavalactones revealed clear SAR, with DMY exhibiting the strongest inhibition of LPS-induced PGE₂ while DHK revealed minimal effects despite their high structural similarity, opening opportunities for future systematic medicine chemistry campaign. Notably, the LPS concentrations used for screening spanned a broad range, with the two lowest doses within or close to the clinically relevant endotoxin range, whereas the higher doses were included as supraphysiologic conditions to elicit robust inflammatory responses *in vitro*. This design allowed anti-inflammatory activity to be evaluated under multiple levels of inflammatory stimulation, thereby increasing the robustness and translational potential of the screening results. Mechanistic elucidation showed that DMY attenuated COX-2 induction in parallel with reduced CREB phosphorylation, indicating CREB-dependent transcriptional control of COX-2 in inflammatory settings, which were further supported by pharmacologic inhibition of PKA. Similarly, CSC induced IL-6, TNF-α and PGE_2_ release in RAW 264.7 macrophages; DMY significantly suppressed these inflammatory responses while DHK had minimal effects. Notably, DMY also reduced CSC-induced p-CREB and COX-2 instead of DHK, providing a common mechanistic signaling between the LPS- and CSC-induced inflammation and AB-free kava and kavalactone suppression. Importantly, the same CREB/COX-2 cascade was aligned with *in vivo* preclinical observations, where cigarette smoke elevated lung p-CREB and COX-2, and AB-free kava and kavalactone dietary supplementation attenuated these changes, connecting the cellular mechanism to cigarette smoke-associated lung inflammation in a preclinical model. Our data also suggests that the anti-inflammatory effects of AB-free kava and kavalactones were independent of NF-κB and AP-1 signaling.

A notable outcome is the divergence between *in vitro* potency and *in vivo* efficacy rankings among kavalactones. Although DMY displayed the highest intrinsic potency in suppressing LPS-induced PGE₂ in macrophages, the cigarette smoke-exposed mouse model showed the most robust anti-inflammatory effects with AB-free kava and DHM, followed by DMY, with DHK comparatively ineffective, including neutrophil accumulation and elevation of TNF-α, IL-6 and PGE_2_. DHM also outperformed DMY *in vivo* in suppressing cigarette smoke-induced p-CREB and COX-2 induction. This divergence (cellular *in vitro* potency does not necessarily translate into whole-animal *in vivo* efficacy) is likely driven by their distinct differences in *in vivo* bioavailability. We propose that differential bioavailability and/or local lung concentration contributes to the superior *in vivo* performance of DHM relative to DMY, despite DMY’s higher *in vitro* potency. At the same time, AB-free kava delivers multiple constituents simultaneously, leading to the possibility of additive or synergistic anti-inflammatory effects and/or differential tissue distributions that may enhance effective exposure and scope of key bioactive components *in vivo*, which requires future more systematic investigation to prioritize the top candidate(s) for translational development.

These findings have several limitations that require future investigations. First, given the challenges of modeling cigarette smoke-driven PGE₂ induction *in vitro* and *in vivo*, quantification of more stable downstream metabolites (e.g., PGEM) and broader lipid mediator profiling may better capture smoke-associated remodeling. Second, PK–PD integration will be critical to test the proposed exposure-driven explanation for *in vivo* ranking by quantifying plasma and lung levels of DHM, DMY, and DHK during dietary supplementation and relating exposure to BAL cytokines, PGE₂, and lung signaling endpoints. Finally, extension of key findings to primary macrophages and additional lung-relevant cell types will further enhance translational relevance.

In summary, stimulated by the anti-inflammatory potential of AB-free kava in a cigarette smoke–exposed mouse model, we identify bioactive kavalactones that suppress cigarette smoke- and LPS-triggered lung inflammation and elucidate a mechanistic axis linking kavalactone treatment to reduce PKA-dependent CREB phosphorylation, decrease COX-2 induction, and reduce PGE₂ production. While LPS is not a fully faithful surrogate for cigarette smoke exposure-associated inflammation, it provided a robust platform for constituent screening, ranking and mechanistic elucidation. Coupled with CSC-based inflammatory suppression and *in vivo* validation, these data highlight AB-free kava and selected kavalactones (DHM and DMY) as promising anti-inflammatory interventions for cigarette smoke-associated lung inflammation with the PKA/CREB/COX-2 signaling pathway as the underlying mechanism.

## Supplementary Materials

The following supporting information can be downloaded: Figure S1. Calibration curve for PGE₂ quantification by UPLC–MS/MS; Figure S2. Dose- and time-dependent induction of PGE₂ production by LPS in RAW 264.7 macrophages; Table S1. Antibodies used for Western blotting analyses in this study.

## Abbreviations

The following abbreviations are used in this manuscript

AP-1: Activator protein 1
BAL: Bronchoalveolar lavage
BSA: Bovine serum albumin
COPD: Chronic obstructive pulmonary disease
CREB: cAMP response element-binding protein
CSC: Cigarette smoke condensate
DHK: Dihydrokavain
DHM: Dihydromethysticin
DMY: Desmethoxyyangonin
K: Kavain
M: Methysticin
Y: Yangonin
IL-6: Interleukin-6
LPS: Lipopolysaccharide
NF-κB: Nuclear factor kappa B
PGE₂: Prostaglandin E₂
PKA: Protein kinase A
RAW 264.7: RAW 264.7 murine macrophages
TLR4: Toll-like receptor 4
CS: Cigarette smoke
TNF-α: Tumor necrosis factor alpha

