## supplemental figure 1&2 Table 1 for "AB-Free Kava and its Kavalactones Suppress Cigarette Smoke- and Lipopolysaccharide-Induced Lung Inflammation: Efficacy and Mechanisms"

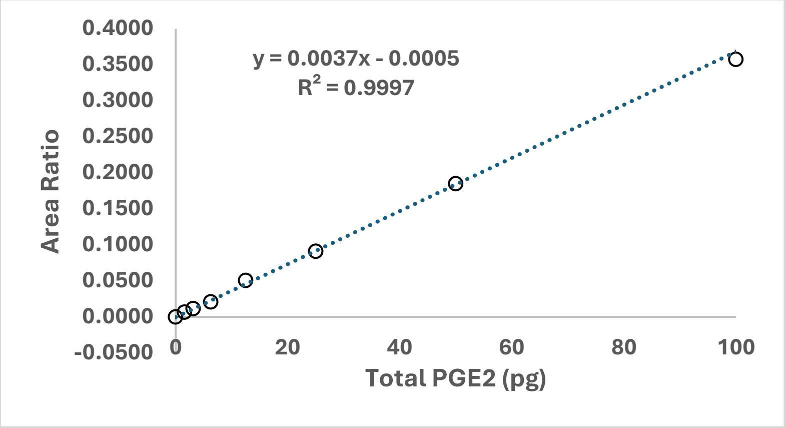


**Figure S1. Calibration curve for $PGE_2$ quantification by UPLC–MS/MS.**

An eight-point calibration curve was prepared by spiking $PGE_2$ standards into cell culture medium over the range of 1.5625-200 pg on column. The limit of detection (LOD) and limit of quantification (LOQ) were estimated as 1.69 and 5.14 pg on column, respectively, based on 3.3σ/s and 10σ/s, where σ represents the standard deviation of the response and s is the slope of the calibration curve.

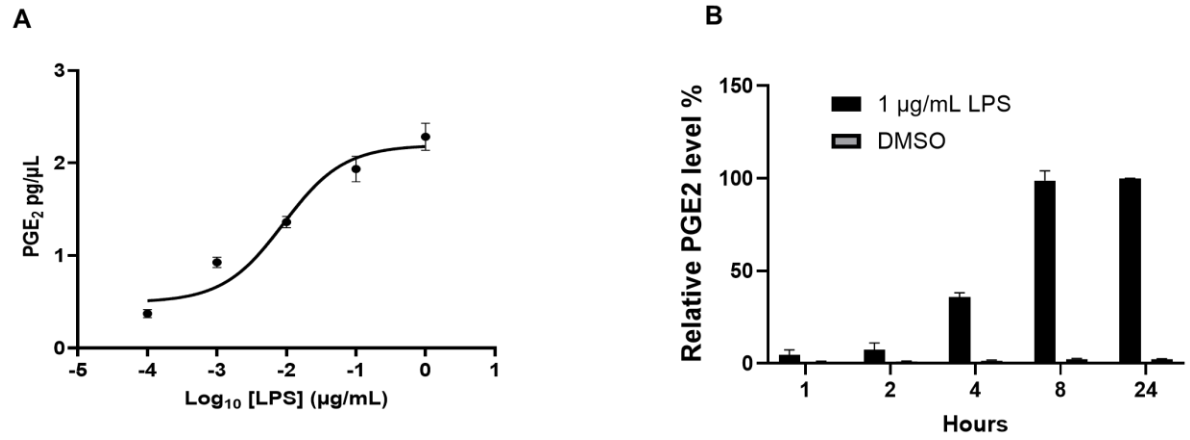


**Fig. S2**. **LPS stimulation robustly induces $PGE_2$ production in RAW 264.7 macrophages in a dose- and time-dependent manner.** (A) Dose–response analysis of LPS-induced $PGE_2$ production. Cells were treated with increasing concentrations of LPS for 8 h, and $PGE_2$ levels in culture supernatants were quantified. LPS stimulation resulted in a clear concentration-dependent increase in $PGE_2$ production, reaching a plateau at 1 µg/mL. (B) Time-course analysis of $PGE_2$ production following LPS stimulation (1 µg/mL). $PGE_2$ levels were measured at the indicated time points. $PGE_2$ production increased progressively and reached a peak at approximately 8 h post-stimulation. Data are presented as mean ± SD.

Table S1. Information of Antibodies used for Western Blotting analyses in this study

| Antibodies | Company | Catalog number |
|---|---|---|
| CREB (48H2) Rabbit mAb | Cell Signaling Technology | 9197 |
| Phospho-CREB (Ser133) (87G3) Rabbit mAb | Cell Signaling Technology | 9198 |
| P-c-Jun (S63) (54B3) Rabbit mAb | Cell Signaling Technology | 2361P |
| P-NF-kappaB p65 (S536) (93H1) Rabbit mAb | Cell Signaling Technology | 3033S |
| Cox2 (D5H5) XP(R) Rabbit mAb | Cell Signaling Technology | 12282 |
| Anti-rabbit IgG, HRP-linked Antibody | Cell Signaling Technology | 7074 |
| Mouse monoclonal anti-beta-actin | Sigma Aldrich | A2228 |
